# Cholesterol-Dependent Dimerization and Conformational Dynamics of EphA2 Receptors: Insights from Coarse-Grained and All-Atom Simulations

**DOI:** 10.1101/2025.01.07.631553

**Authors:** Amita Rani Sahoo, Nisha Bhattarai, Matthias Buck

## Abstract

The EphA2 transmembrane receptor regulates cellular growth, differentiation, and motility, and its overexpression in various cancers makes it a potential biomarker for clinical cancer management. EphA2 signaling occurs through ligand-induced dimerization, where the transmembrane (TM) and juxtamembrane (JM) domains play crucial roles in stabilizing the dimer conformations and thereby facilitating signal transduction. Electrostatic interactions between basic JM residues and signaling lipids (PIP2 and PIP3) regulate phosphorylation while Cholesterol’s potential role in modulating EphA2 activation remains unclear. To investigate this, we modeled the TM-full JM peptide of EphA2 and employed coarse-grain and all-atom simulations to investigate its dimerization in cholesterol-rich and cholesterol-deficient membranes. Our findings reveal that cholesterol stabilizes specific TM dimers and TM-JM interactions with PIP2, highlighting the importance of membrane composition in EphA2 dimerization, oligomerization, and clustering. These insights enhance our understanding of lipid-mediated regulation of EphA2 and its implications in receptor signaling and cancer progression.

## Introduction

Ephrin receptors (Ephs) are key members of the receptor tyrosine kinase (RTK) family, playing essential roles in developmental processes and contributing to the progression of various cancers.^1–3^ Typically, Eph receptor activation is triggered by interactions with membrane-anchored ephrin ligands. These ligand-receptor interactions lead to receptor dimerization, activation of the cytoplasmic tyrosine kinase domain, followed in the case of EphA2 by the formation of higher-order oligomeric clusters.^4^ Posttranslational modifications also enable signaling through phosphotyrosine-mediated interactions.

Structurally, Eph receptors^5,6^ consist of a large N-terminal extracellular region containing the ligand-binding domain (LBD), a cysteine-rich domain (CRD), and two fibronectin type III (FNIII) domains. This extracellular region connects to the intracellular segment via a single transmembrane (TM) domain. The intracellular region includes a flexible juxtamembrane (JM) region, a tyrosine kinase domain (KD), a sterile-alpha motif (SAM), as well as a PDZ binding motif (Figure 1A). Interestingly, Eph receptors can dimerize independently of ligand binding under conditions of high receptor concentration within the plasma membrane. This ligand-independent dimerization has been shown to lead to a different, non-canonical function, driven not by kinase function but by their role as substrates for serine/threonine phosphorylation by various S/T kinases.^7,8^ Thus, dimerization and oligomerization, whether ligand-dependent or independent, can result in distinct signaling outcomes. For instance, EphA2 can act as an oncogene through ligand-independent mechanisms while serving as a tumor suppressor that facilitates cell repulsion and positional stability when its tyrosine kinase domain is activated by ligand-induced clustering.^9,10^

**Figure 1.**
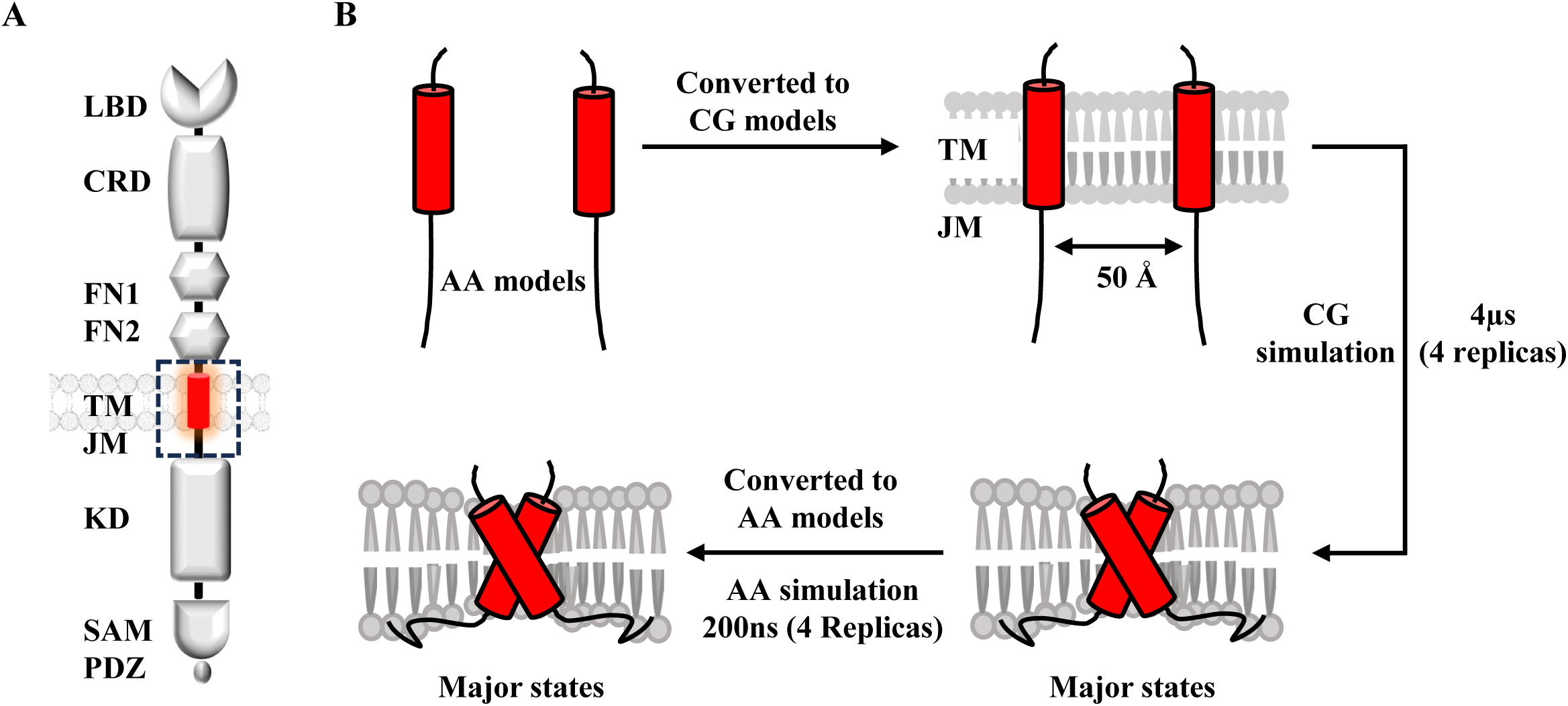
(A) Schematic representation of the EphA2 receptor. The extracellular region comprises the ligand-binding domain (LBD), a cysteine-rich EGF-like domain (CRD), and two fibronectin type III repeats (FN1, FN2), followed by the transmembrane (TM) domain. The intracellular region includes the juxtamembrane (JM) domain, tyrosine kinase domain (KD), sterile alpha motif (SAM), and PDZ-binding motif (PDZ). (B) Overview of the study methodology. All-atom (AA) monomeric peptide models were first converted to coarse-grain (CG) representations and embedded in a lipid bilayer for CG simulations. The dominant conformers from CG simulations were then back-mapped to AA models and subjected to AA simulations for further analysis.

While structural studies of isolated RTK domains, such as the ectodomain,^5,11^ transmembrane (TM) domain,^12^ kinase and SAM domains,^7,13,14^ have provided significant insights, integration of these distinct domains into a comprehensive model for the receptor’s function within the membrane has remained challenging. Furthermore, the interplay between Eph receptors and membrane lipids, such as glycolipids and phosphatidylinositol phosphates (PIPs), must be considered to fully understand receptor function.^15^ Cholesterol, which constitutes approximately 30% of the lipid bilayer of the cell’s plasma membrane, plays a vital role in maintaining membrane integrity and regulating fluidity, significantly influencing membrane protein dynamics.^16,17^ Specifically, cholesterol content has been shown to affect the rigidity and fluidity of plasma membranes, impacting receptor behavior. A study by Zhang et al.^18^ demonstrated that cholesterol modulates ErbB2 receptor dynamics, promoting endocytic degradation in breast cancer cells when cholesterol levels are reduced. However, the specific effects of cholesterol on Eph receptor signaling remain unclear. Our recent collaborative study with the Barrera lab.^17^ explored how cholesterol influences EphA2 self-assembly and activity, emphasizing its potential role in modulating receptor function. The study revealed that cholesterol inhibits EphA2 activation through an in-trans mechanism involving protein kinase A, regulated by beta-adrenergic receptor signaling. However, our previous simulations focused only on the transmembrane (TM) region and a short juxtamembrane (JM) segment.

The rapid advancements in molecular dynamics (MD) simulations have enabled detailed investigations of membrane proteins and of their interactions with lipids. These simulations have been particularly valuable in studying RTKs such as EGFR and related receptors.^19,20^ In this study, we applied both coarse-grained (CG) and all-atom (AA) molecular dynamics simulations to understand the dimerization mechanism of the EphA2 receptor in the absence of its ligand. Specifically, we examined the transmembrane (TM) and juxtamembrane (JM) domains to uncover their roles in receptor association and function. Our study focused on modeling the full TM and JM regions (residues S^528^–C^612^) of human EphA2 and analyzing their dimerization within cholesterol-rich and cholesterol-deficient membrane environments. We observed distinct conformational transitions in EphA2 dimers depending on membrane cholesterol levels: between left-handed (inactive) and right-handed (active) dimers in cholesterol-rich membranes and between left-handed and parallel dimers in cholesterol-deficient membranes. However, the NMR structure^12^ showed a left-handed configuration with a crossing angle of 15° and contacts formed via an extended heptad motif (^535^LxxxGxxAxxxVxx^549^L). Although the TM sequence also contains a glycine zipper motif (^536^AxxxGxxx^544^G) compatible with a -45° right-handed structure predicted by MD simulations, this was not observed experimentally. However, point mutations and kinase assays confirmed the switch between heptad and glycine zipper-stabilized structures, depending on ligand presence.^21^ Extensive simulations with an earlier Martini force field reproduced a near-parallel configuration (crossing angle ∼10°) with heptad-like contacts but also transitioned to a right-handed configuration (-20° crossing angle).^22^ In our previous study using Martini 3 in a POPC bilayer, the right-handed glycine zipper structure was predominant in presence of the activator peptide and was associated with ligand-independent receptor activation.^23^ Our study demonstrates that the dimerization and conformational transitions of the EphA2 TM and JM regions are modulated by membrane composition, with cholesterol stabilizing distinct active and inactive dimer states and interactions with anionic lipids, such as PIP2, influencing receptor activation and membrane dynamics. These findings provide novel insights into the interplay between lipid composition and EphA2 receptor dimerization, contributing to our understanding of its ligand-independent signaling mechanisms.

## Results

### Association of TM-full JM peptides

For our investigation into the association of EphA2 TM-full JM peptides, we harnessed the power of Coarse Grain (CG) molecular simulations. As our initial setup, as shown in Figure 1B, we modeled the TM-full JM peptide with the TM region extracted from the NMR structure and the JM region as an extended conformation. Subsequently, we set up two identical TM-full JM peptides, positioning them parallel to the membrane normal and at the lipid bilayer’s core, ensuring a gap of 50Å between them. It is well established that the configuration of TM helices can be influenced not only by membrane thickness but also by lipid composition, including factors such as the presence of cholesterol. This understanding guided our strategy in constructing two distinct systems, each with different membrane compositions. Specifically, system 1 embraced a combination of POPC (55%), Cholesterol (40%), and PIP2 (5%). In contrast, system 2 was structured with an alternate composition, predominantly comprising POPC (95%) along with PIP2 (5%). We conducted the CG simulation over a span of 4 μs, running six replicas for each system to ensure result consistency. To track the association dynamics of the TM-full JM peptides, we observed the distance between the TMs’ center of mass (COM), as illustrated in Figures 2A & 2B. For system 1, which includes cholesterol (Figure 2A), the two helices approached each other within 0.5-1.5 µs in four simulations. By comparison, for system 2, which lacks cholesterol (Figure 2B), the association between the helices was faster, taking approximately 0.5-1 µs in four out of six simulations. Regardless, in both scenarios, the TM-full JM peptides formed dimers and maintained stability throughout the remainder of the simulation period. To streamline the analysis across both systems, we merged the results from all six repeat simulations. We then computed the total populations of the TM dimer configurations, considering both the inter-helical angle and distance between the monomers. These results are visualized in a two-dimensional (2D) plot, as depicted in Figures 2C & 2D. In the cholesterol-containing system (Figure 2C), the 2D plot reveals two predominant global minima which including left-handed TM dimer configurations (shown as 1^st^ Cluster) and right-handed TM dimer configurations (shown as 2^nd^ Cluster). In contrast, the system devoid of cholesterol also presents two prominent global minima, as highlighted in Figure 2D suggesting the existence of left-handed TM dimer configurations (shown as 1^st^ Cluster) and nearly parallel TM dimer configurations (shown as 2^nd^ Cluster). We also noted conformational transitions in TM dimers at varying crossing angles across both systems. Additionally, we examined whether the intermolecular interactions among TM dimers exhibited differences between the two systems. To investigate this, we computed the average contact map across residue interfaces within the TM regions, as displayed in Figure 3. Remarkably, in the presence of cholesterol, both left- and right-handed TM dimers predominantly utilized the heptad repeat motif (Figure 3A) interface. Conversely, in the absence of cholesterol, we observed involvement of both the GxxxG motif (influenced by the parallel TM dimers) and the heptad repeat motif within the contact interface (Figure 3B).

**Figure 2.**
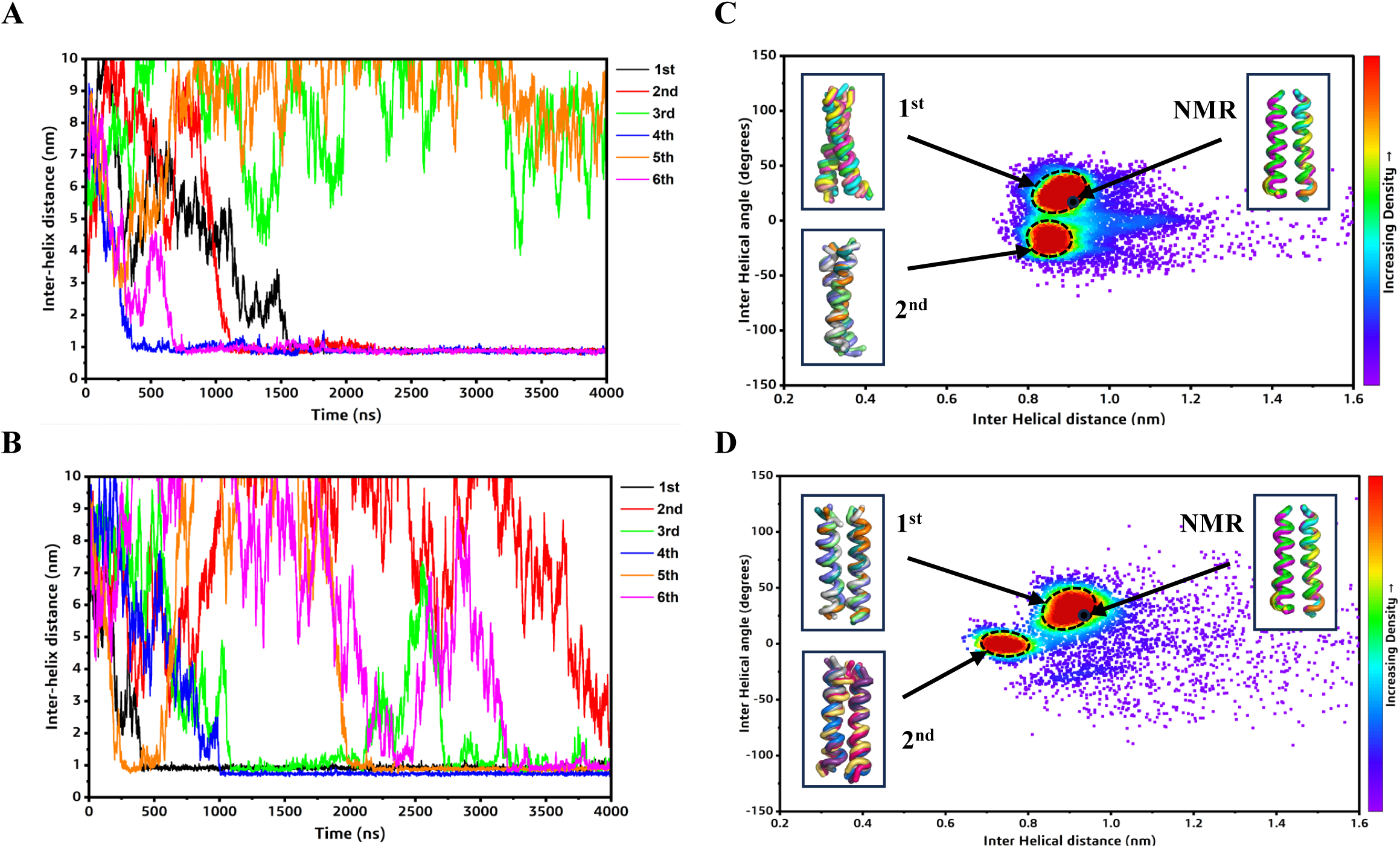
Comparison of the association of the EphA2 TM regions from the CG simulations. Left panel-Inter-helical distance (COM) plots showing the association between the TM regions of EphA2 in presence (A) and absence of cholesterol (B) in the membrane. 1st, 2nd, 3rd, 4th, 5th and 6th simulation results are shown in black, red, green, blue, orange and magenta lines, respectively. Right panel-2D distribution plot (interhelix angle vs. distance) between the EphA2 TMs in the presence (C) and absence of cholesterol (D) in the membrane. Data from the last 2 µs of the simulations are used for 2D plot analysis. Coordinates for the NMR structure are marked as black circle. Representative poses from each population are displayed as multicolored ribbon. All the replica trajectories are considered for analysis.

**Figure 3.**
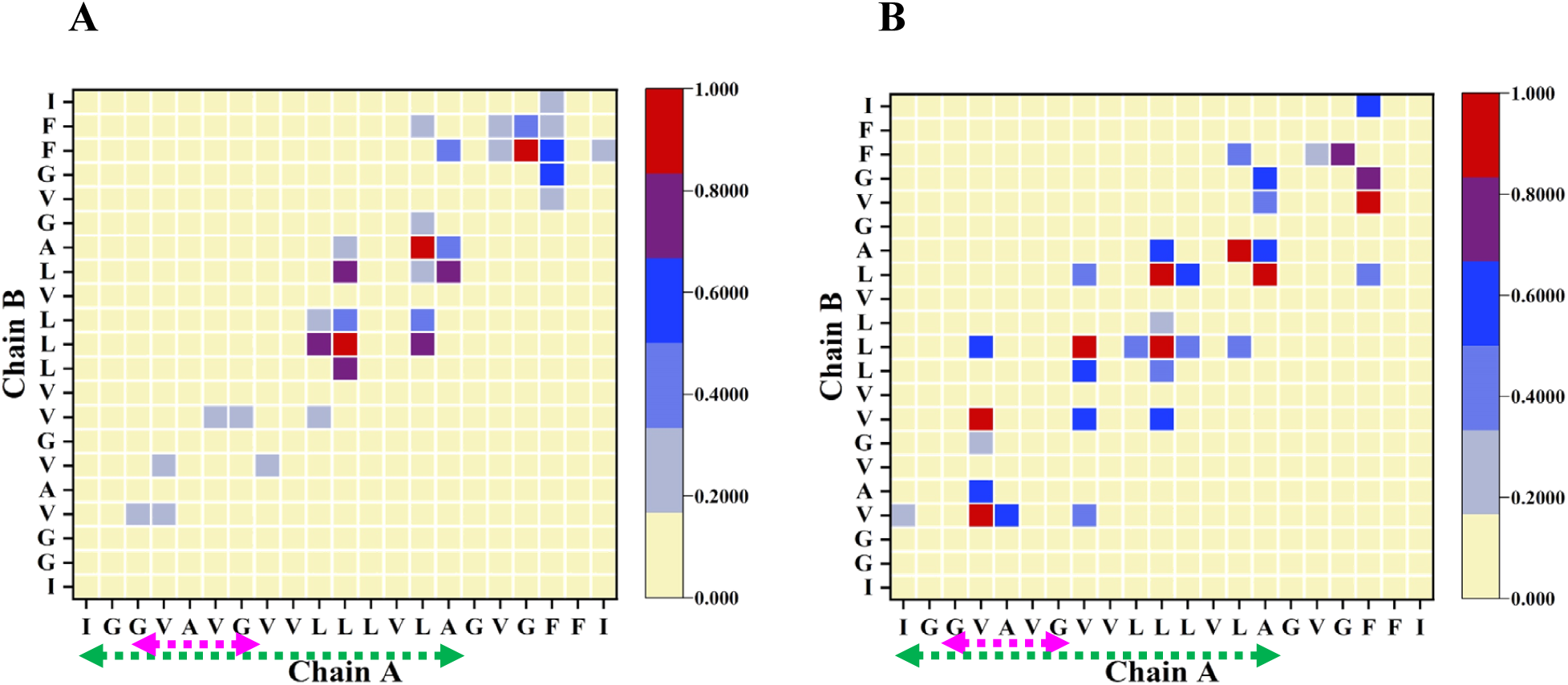
Comparison of the average contact map interface between the EphA2 TMs in the presence (A) and absence (B) of Cholesterol in the membrane. Contact maps are calculated with a 6Å cut-off. The color scale (yellow to blue to red) indicates the fractional occupation of TM contacts (0 to 1). Data from the last 2 µs from all the replica trajectories of the CG simulations are used for analysis. Heptad and GxxxG motifs are labeled as green and pink arrows.

### Involvement of the CRAC motif for cholesterol binding

To further investigate the role of cholesterol, we analyzed the interactions between EphA2 TM-full JM peptides with cholesterol as presented in Figure S1. We tracked the average count of interactions involving cholesterol molecules for each amino acid in the sequence of the TM-JM peptide, within a cut-off of 0.6 nm, covering the entirety of the simulation period. It is noteworthy that most interactions of cholesterol with the TM region remained stable over the course of the simulation. Of particular interest are also F556 and F557 which are components of a putative Cholesterol Recognition/interaction Amino acid Consensus (CRAC) domain.^24^ These exhibited remarkably strong interactions with cholesterol molecules. However, CRAC domains have a sequence motif of (V-X_1-5_-F-X_1-5_-R) and interactions with cholesterol only involve the N-terminal and middle part of this sequence and few JM residues near the C-terminus displayed some interactions with cholesterol. Furthermore, the radial distribution function of all lipids around the protein was computed for both systems, illustrated in Figure S2. Interestingly, PIP2 exhibited a particularly strong interaction with the protein, displaying a more pronounced affinity than cholesterol. There is a slight, but likely not statistically significant increase in PIP2 around the protein in the presence of cholesterol, while cholesterol is more excluded from the proximity of the protein on average compared to POPC lipid.

Further, we extracted few representative TM dimer structures from each population cluster (Figures 2C & 2D) from both the systems and compared with the NMR structure. Based on the best compared RMSD with the NMR structure (table S1), we selected one TM-full JM dimer conformation from each population for both the systems and subjected to all-atom simulations for 200ns each to get more detailed atomistic insights.

### Detailed Interactions of dimer with Cholesterol and PIP_2_ (system1)

With slow conformational dynamics within the lipid bilayer, the two distinct sub-systems in the cholesterol-rich environment of system 1—originating from left- and right-handed dimer conformations—exhibited different interaction patterns with the lipid bilayer. We clustered the dimers from both sub-systems using a backbone RMSD of 1.5 nm to identify the major conformational clusters. Representative structures of the dominant population clusters for both the left- and right-handed dimers are depicted in Figure 4.

**Figure 4.**
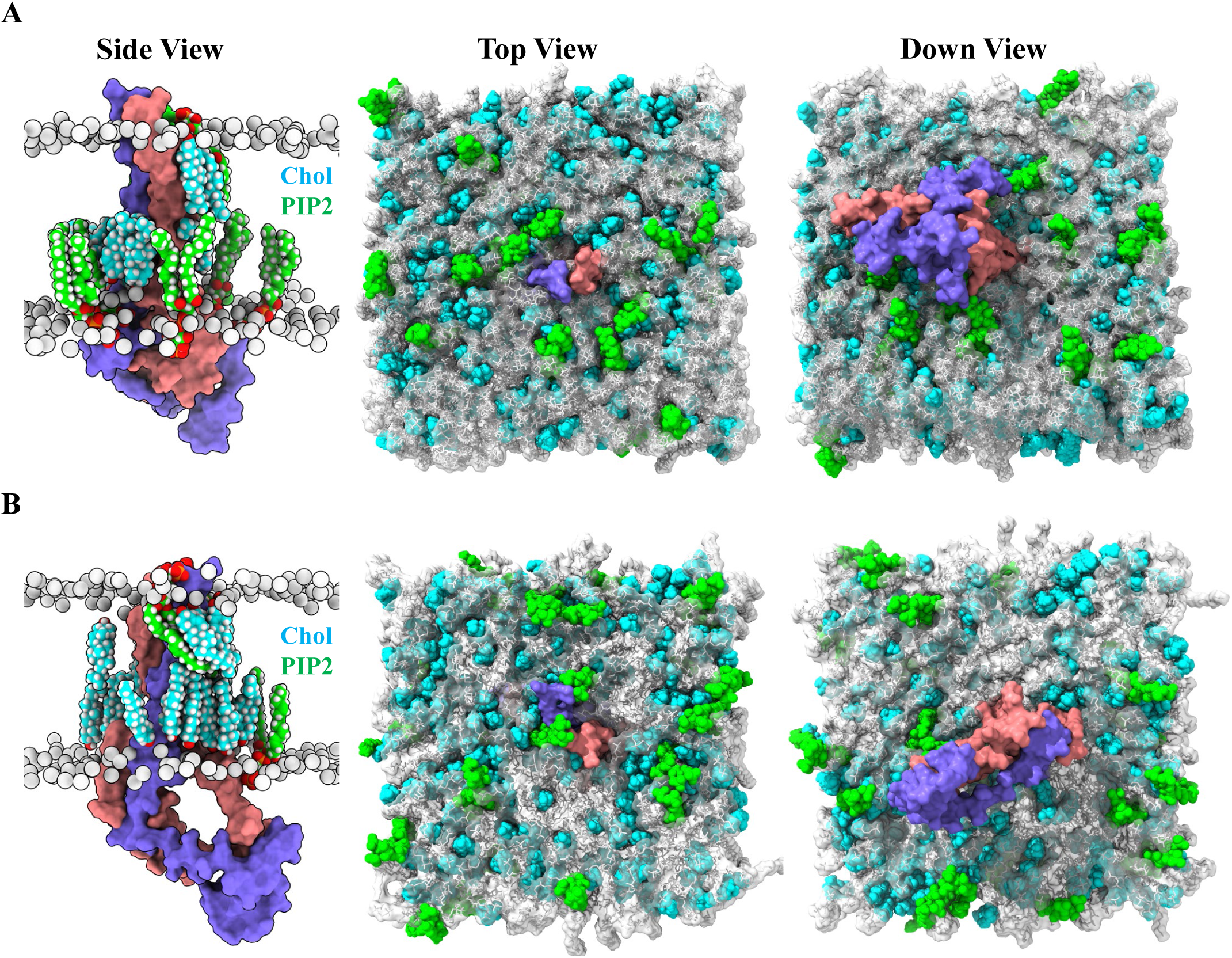
Comparative snapshots of representatives of major population clusters from cholesterol rich systems from the AA simulations. Different views of the left-handed (A) and the right-handed (B) dimer surrounded by lipids. EphA2 TM-full JM peptides are shown in orange red and purple surfaces. Cholesterol and PIP2 are shown in cyan and green spheres. POPC are shown as light grey sticks with surface transparency of 60%.

We analyzed the cholesterol occupancy in both structures, also referred to as configurations, and found that cholesterol molecules strongly interacted with the TM regions in both cases (Figure 5), which partially aligns with the location of the CRAC domain (see above) and is consistent with its known role in cholesterol interaction. Additionally, we examined the binding poses of cholesterol with the TM-full JM dimer, revealing that the predominant interactions occur with the TM regions (site1-site4 for left-handed and site1-site3 for right-handed) rather than with the JM regions.

**Figure 5.**
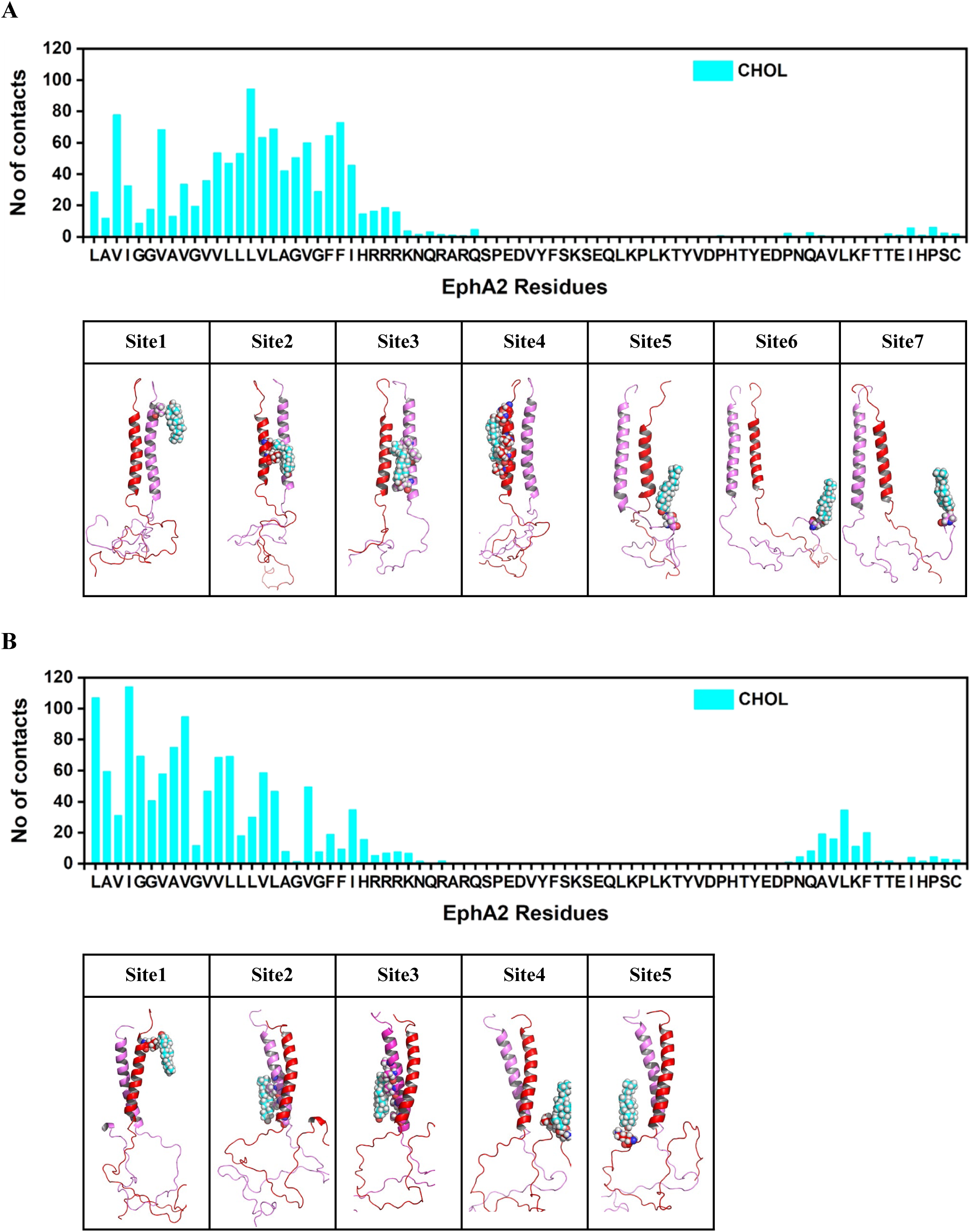
Comparative occupancy of cholesterol-protein interaction of left-handed (A) vs right-handed (B) configurations from the AA simulations. Upper panels represent residue specific interaction of cholesterol and lower panels represent different binding sites/ poses of cholesterol molecules with the dimer. The interacting cholesterol molecules and the binding site atoms of the protein are shown as spheres. We used 0.7 nm cutoff distance for contact calculation.

Similarly, we compared the PIP2 occupancy across both structures and observed distinct interaction patterns between the left- and right-handed configurations (Figure 6). In the left-handed configuration, PIP2 lipids primarily occupy the JM region, as expected, whereas in the right-handed configuration, PIP2 lipids interact more with the TM regions compared to the JM region. As illustrated, the PIP2 binding poses for the left-handed structure are more prevalent in the JM region (sites 2-4) than in the TM region (site 1). Conversely, in the right-handed structure, PIP2 binding poses are more frequent in the TM region (sites 1-2) than in the JM region (sites 3-4).

**Figure 6.**
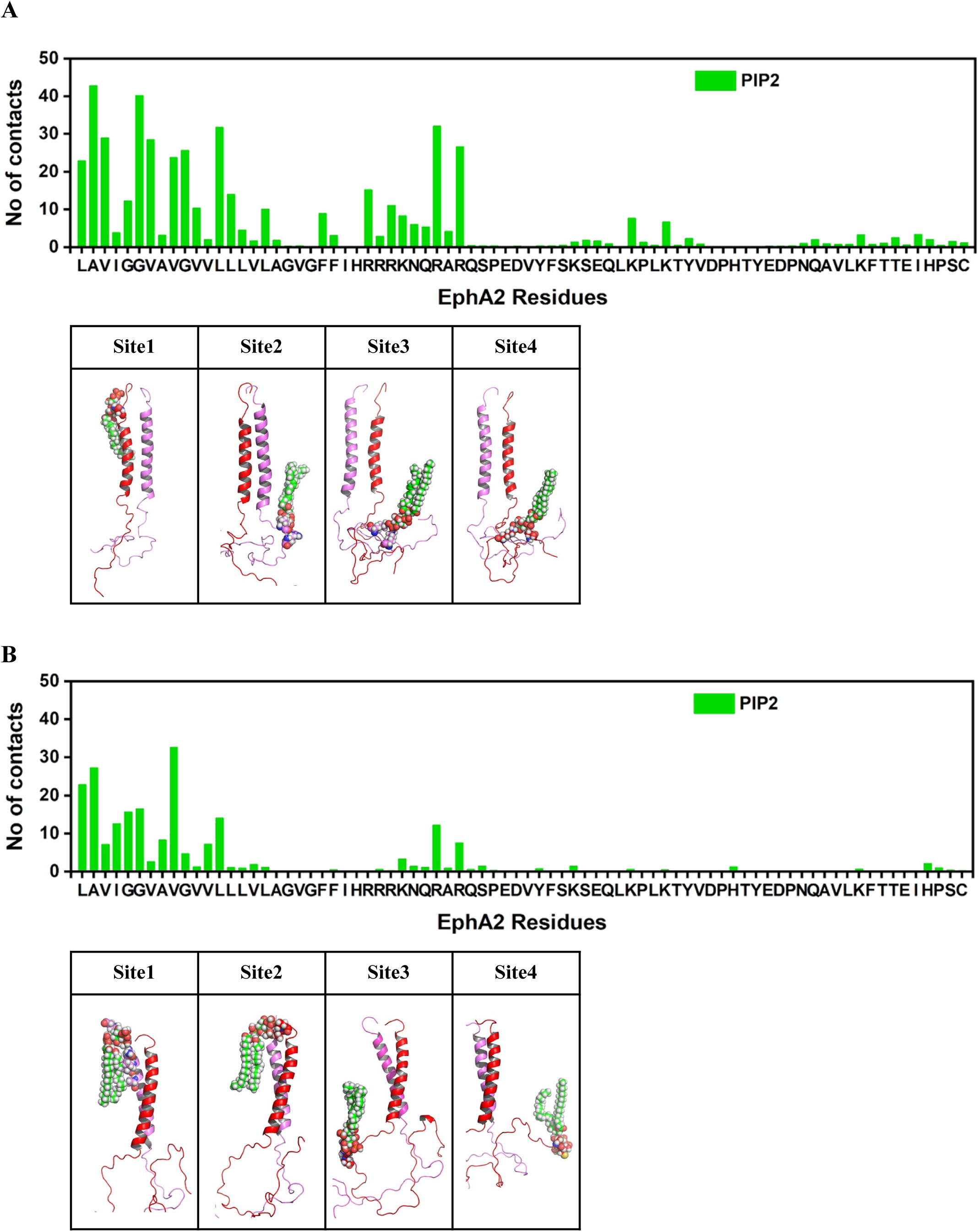
Comparative occupancy of PIP2-protein interaction of left-handed (A) vs right-handed (B) configurations from the AA simulations in cholesterol-rich membrane systems. Upper panels represent residue specific interaction of PIP2 and lower panels represent different binding sites/ poses of PIP2 molecules with the dimer. The interacting PIP2 molecules and the binding site atoms of the protein are shown as spheres. We used 0.7 nm cutoff distance for contact calculation.

### Interaction of dimer with PIP_2_ in absence of cholesterol (system2)

In the cholesterol-deficient system 2, the two distinct configurations, which began from left-handed and parallel TM dimer conformations, exhibit different binding patterns of PIP2 lipids. We clustered the dimers from both configurations using a backbone RMSD of 1.5 nm to identify the major conformational clusters. Representative structures of the dominant population clusters for both the left-handed and parallel dimers are shown in Figure 7.

**Figure 7.**
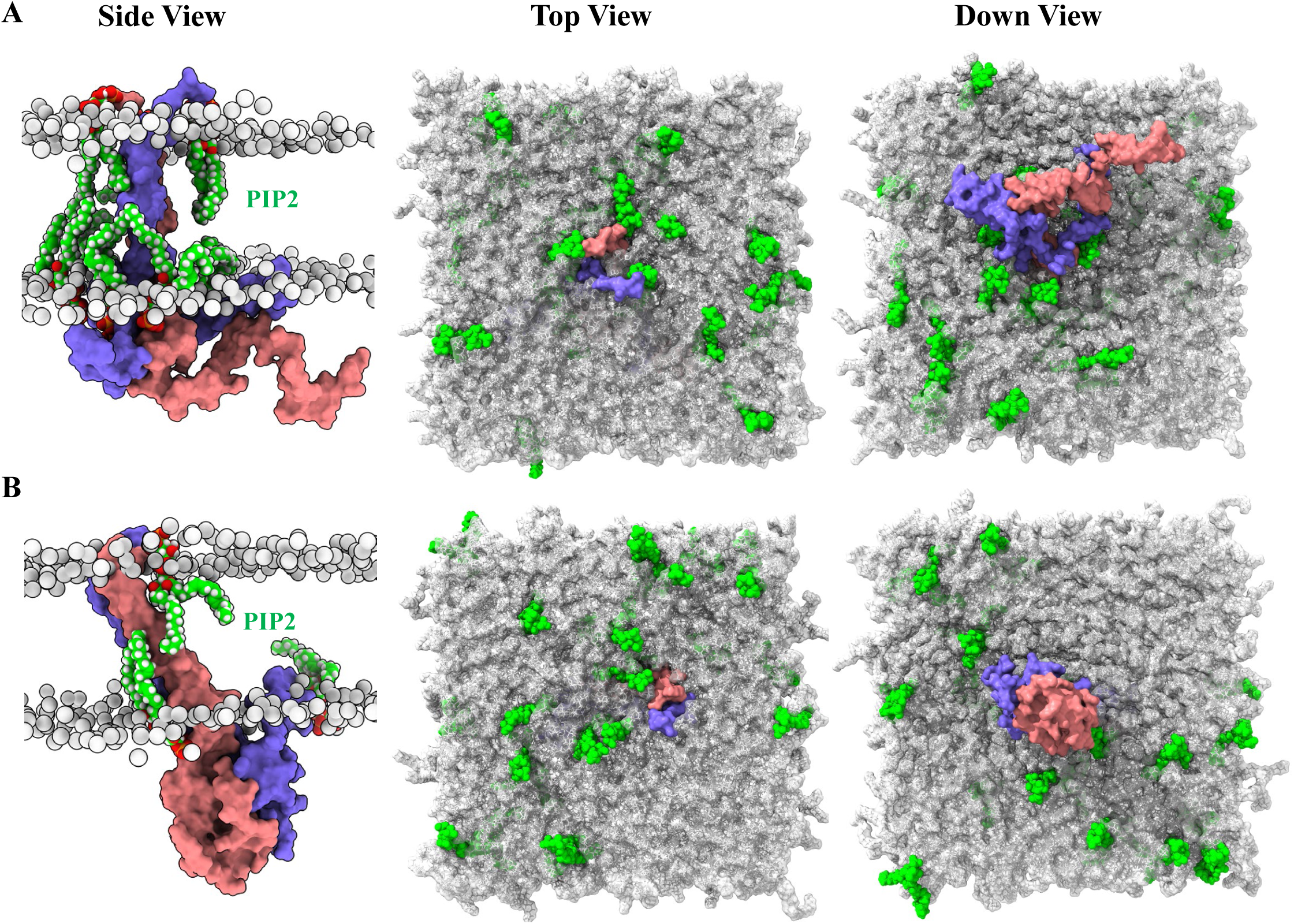
Comparative snapshots of representatives of major population clusters from cholesterol deficient systems from the AA simulations. Different views of the left-handed (A) and the parallel (B) dimer surrounded by lipids. EphA2 TM-full JM peptides are shown in orange red and purple surfaces. PIP2 molecules are shown as green spheres. POPC are shown as light grey sticks with surface transparency of 60%.

In the absence of cholesterol, PIP2 lipids interact across both the TM and JM regions in the left-handed structure. However, in the parallel dimer, PIP2 lipids primarily occupy the latter part of the TM region and the beginning of the JM region (Figure 8). For the left-handed structure, PIP2 binding poses are most prevalent at sites 1, 3-4, and 6, with lower occupancy at sites 2 and 5. Conversely, in the parallel dimer, site 2 shows higher prevalence compared to other sites (data not shown).

**Figure 8.**
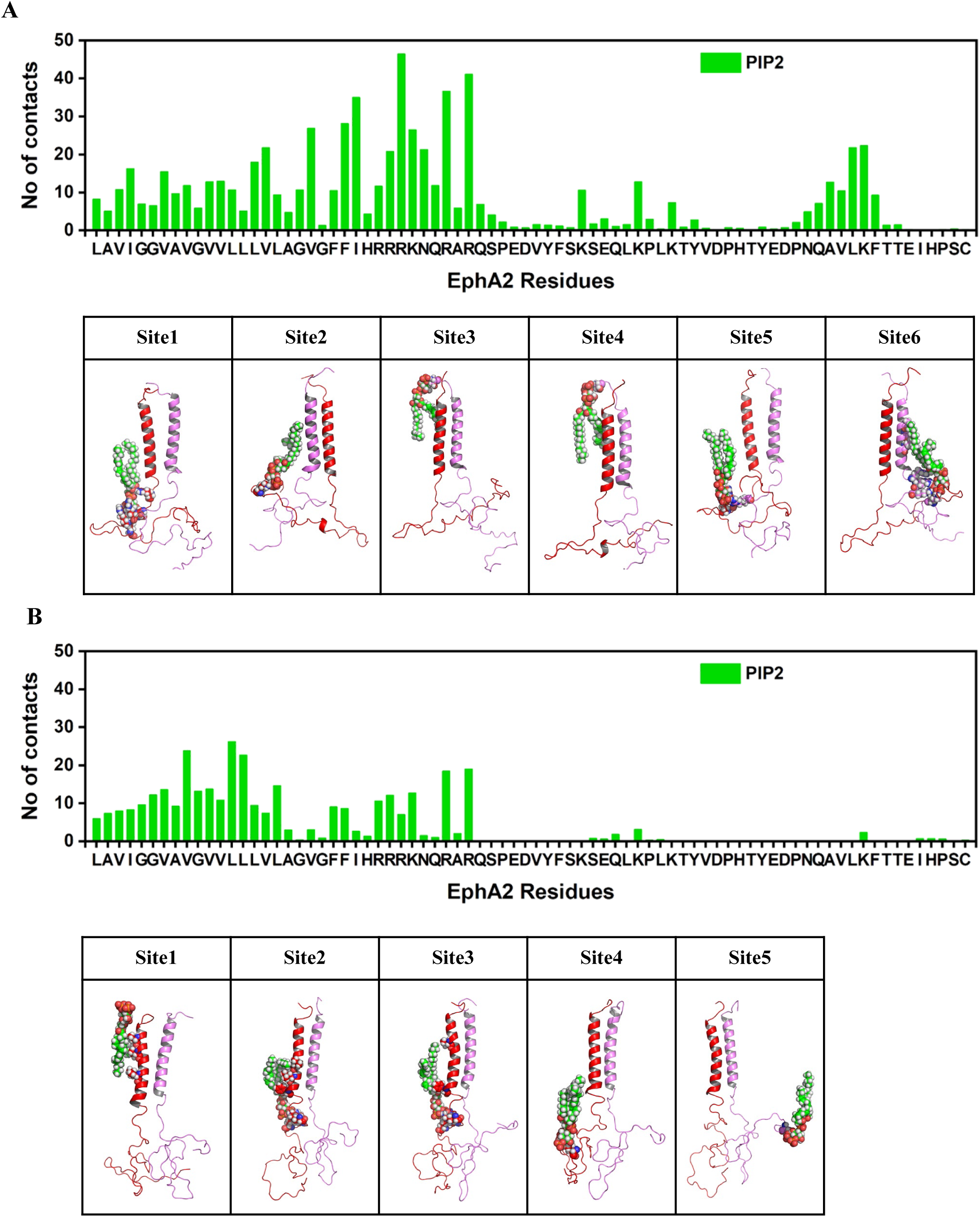
Comparative occupancy of PIP2-protein interaction of left-handed (A) vs parallel (B) configurations from the AA simulations in cholesterol-deficient membrane systems. Upper panels represent residue specific interaction of PIP2 and lower panels represent different binding sites/ poses of PIP2 molecules with the dimer. We used 0.7 nm cutoff distance for contact calculation.

### Overall effect of membrane packing in helix association

A visual inspection of the trajectories revealed significant difference in membrane curvature surrounding the dimers in both cholesterol-rich and cholesterol-deficient environments (Figure 9). To characterize the local membrane perturbations, we calculated the differences in the membrane packing around the dimerized states (left and right-handed dimer) for the cholesterol-rich systems and (left-handed and parallel dimers) for the cholesterol-deficient systems. The bilayer thickness was calculated for the entire bilayer by binning into 0.1nm^2^ grids and is shown in Figure S3. Large variations were observed in the bilayer thickness of the lipids around each of the dimer, consistent with the curvature analysis above. In cholesterol-rich membranes (Figures S3A & B), a reduction in bilayer thickness (giving a thickness of ∼1.5 to 2.0 nm) was observed surrounding both the left- and right-handed dimers, followed by a broader increase in bilayer thickness (to a thickness of ∼3 to 5 nm). Conversely, in cholesterol-deficient membranes (Figures S3C & D), there was an increase in bilayer thickness (total thickness ∼2.0 to 2.8 nm) around both the left-handed and parallel dimers, with a more pronounced effect in the latter compared to the overall bilayer thickness (thickness ∼3.3 to 4.0 nm). The increased thickness in cholesterol-rich membranes is attributed to cholesterol’s rigid steroid ring structure, which contributes to overall bilayer thickness by interacting with phospholipid acyl chains, modifying lipid packing, and enhancing membrane properties.^25–27^

**Figure 9.**
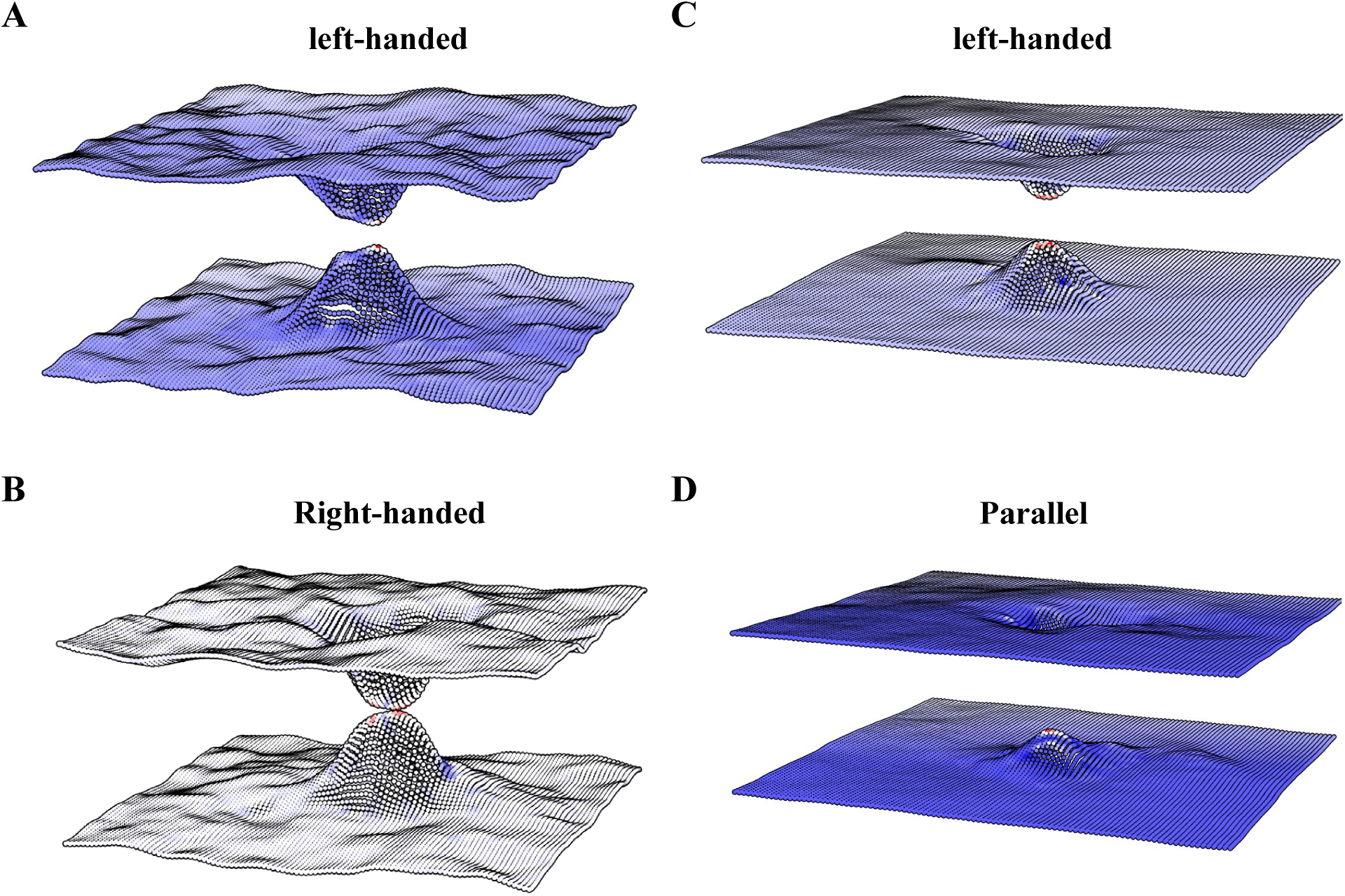
Comparison of membrane curvature calculated over the entire membrane for the Cholesterol rich (A & B) and cholesterol deficient (C & D) systems from the AA simulations. Replica trajectories are combined for this analysis.

We further analyzed the order parameter (SCD) of POPC lipids for all four subsystems (Figure S4). This analysis combined all replica trajectories, providing SCD values for the carbon atoms in both acyl chains (sn-1 and sn-2). The results indicate that cholesterol-containing membranes exhibit higher SCD values compared to cholesterol-deficient systems, reflecting the well-established cholesterol-induced phospholipid ordering effect observed in both experimental and computational studies.^16,28–30^ Cholesterol presence in the membrane appears to enhance rigidity and order relative to cholesterol-deficient systems, integrating into the bilayer and filling spaces between phospholipids, thereby stabilizing the membrane and decreasing fluidity by ordering lipid molecules

## Discussion

Generic but in some cases detailed lipid-protein association determines protein topology, activity, and aggregation.^31^ The association between helical protein segments depends on both specific and non-specific interactions, the latter being mediated by the surrounding lipids and driven by the perturbation of the membrane environment by the protein. Lipid-mediated protein– protein interactions have been extensively studied using theoretical, simulation, and experimental approaches.^32–34^ The association of EphA2 receptors in the absence of the ephrin ligand has been linked to non-canonical signaling pathways.^35,36^ In particular, the role of TM dimerization in the context of the full-length receptor remains unclear. While our previous study demonstrated conformational transitions of the TM peptide within a DMPC bilayer,^37^ the impact of the flexible and unstructured JM region on TM dimerization had not yet been explored. To address this, we employed coarse-grain and all-atom simulations to investigate the association between human EphA2 receptors.

Our findings reveal significant insights into how the lipid environment influences the association and structural dynamics of EphA2 TM-full JM peptides. The interplay between cholesterol, PIP2, and the TM domain highlights critical biophysical principles governing protein dimerization and lipid-protein interactions, with cholesterol helping to resolve hydrophobic mismatches between the protein and membrane bilayer.^38,39^ The results underscore the dual role of cholesterol in modulating both the dynamics and stability of TM dimer configurations. Cholesterol-rich systems favored transitions between left- and right-handed dimers (Figure 2C), suggesting that cholesterol imparts flexibility to the TM dimers while stabilizing specific interaction motifs, such as the heptad repeat (Figure 3A). The stronger cholesterol binding at the CRAC motif ^24^ (Figure S1) highlights its role in stabilizing TM helices through selective interactions. For example, studies on membrane proteins, particularly GPCRs, suggest that cholesterol binding at CRAC motifs plays a key role in stabilizing transmembrane helices, influencing both receptor dimerization and overall receptor function.^40,41^ This cholesterol interaction not only enhances structural stability but also modulates the functional dynamics of membrane proteins, contributing to signaling efficiency and receptor activity. Additionally, membrane packing analyses demonstrate how cholesterol alters bilayer thickness and lipid ordering, enhancing the rigidity of the local environment around dimers. The cholesterol-deficient systems, in contrast, revealed a different dynamic profile. Along with the left-handed TM dimers, parallel TM dimer configurations, facilitated by the GxxxG motif (Figure 3B), emerged as a unique feature in the absence of cholesterol. These dimers demonstrated more pronounced perturbations in membrane thickness and packing, suggesting that cholesterol deficiency might promote less ordered but potentially more flexible membrane environments. Such variations in lipid-induced constraints can significantly influence protein functionality by allowing alternative dimer conformations.

The role of PIP2 was prominent in both systems (Figure S2), but its interaction patterns varied depending on cholesterol presence. Cholesterol-rich membranes exhibited PIP2 binding primarily at TM regions, while in cholesterol-deficient systems, PIP2 interacted across both TM and JM regions (Figures 6 and 8). This observation implies that cholesterol may spatially restrict PIP2 binding, directing it toward regions critical for functional modulation. The specific enrichment of PIP2 near dimer interfaces both in the TM and JM regions also highlights its potential role in stabilizing dimerization and regulating local membrane charge distribution (i.e. clustering of Arg and Lysine sidechains with PIP2), which may influence receptor clustering and signaling as mentioned maintaining the JM region near the membrane, on one hand diminishing its tyrosine phosphorylation, but on the other also moving it away from the kinase domain, relieving its inhibition. Thus, PIP2 is known to promote EphA2dimeriazation and kinase activity.^38^ By contrast, tyrosine phosphorylation of the JM would be expected to move this region away from the membrane, perhaps bringing it again close to the kinase active site for inhibitory interactions. However, it is also possible to have bridging interactions, via cations, which would maintain negatively charged sidechains near PIP2/PS.^42^ The observed differences in bilayer thickness and lipid order parameters (Figures S3 and S4) provide further context for understanding how membrane composition regulates TM peptide association. Cholesterol-rich membranes displayed higher order and thickness due to cholesterol’s rigid structure and its ability to fill gaps between lipid acyl chains. In contrast, cholesterol-deficient membranes exhibited more variable packing and increased fluidity, enabling distinct dimerization behavior and conformational diversity. Importantly, the effect of cholesterol on the membrane thickening would be expected to favor the parallel configuration of helices within the TM dimer (reduced crossing angles)-however the opposite is observed. Our observation of greater membrane indentation and curvature near the TM region in presence of cholesterol resolves this apparent contradiction, as the nearby membrane thickness is actually reduced in presence of cholesterol.

These findings emphasize how the local lipid environment around a TM and JM peptide intricately modulates transmembrane protein behavior by influencing both structure and function. PIP2, with its highly negative charge density, plays a pivotal role in promoting receptor dimerization, potentially by reducing electrostatic repulsion between protein domains. This lipid not only stabilizes the dimeric state but may also exert a regulatory effect on ligand-independent EphA2 activation, similar to its established role in driving dimerization of RTKs such as EGFR.^43,44^ Through such interactions, PIP2 emerges as a critical regulatory element in modulating receptor function and signaling outcomes.

In contrast, cholesterol appears to serve as a negative regulator of EphA2 activation. A recent study of the Barrera lab. shows that its inhibitory effect is mediated through an in-trans mechanism involving protein kinase A (PKA) and beta-adrenergic receptor signaling.^17^ Importantly, studies on C99 dimerization suggest that cholesterol does not directly compete with protein dimerization but enhances the stability of dimeric complexes, highlighting its role in fine-tuning membrane protein interactions.^45^ Additionally, the observed differences in dimerization patterns between systems containing only the TM region ^37^and those incorporating the short^17^ or full JM regions in this study underscore the influence of flanking regions. The flexible juxtamembrane (JM) region appears to modulate TM dimerization and structural stability thereby emphasizing the complexity introduced by these regions in shaping protein behavior within membranes (Table S1).

The differential effects of cholesterol and PIP2 on EphA2 TM-full JM dimerization underscore a lipid-mediated regulatory mechanism critical for receptor activity. These findings underscore the broader significance of membrane composition in shaping receptor signaling, particularly in cancer, where membrane properties are frequently altered. By integrating CG-MD and AT-MD simulations, our study provides valuable insights into lipid-receptor interactions. However, future efforts should extend to simulating the full-length EphA2 dimer within complex lipid mixtures representative of cellular membranes. Importantly, as this study is purely computational, additional research is required to confirm whether JM-PIP2 interactions drive canonical or non-canonical signaling pathways. Combining atomistic simulations with experimental validation will be crucial for unraveling the precise mechanistic impact of lipid-protein interactions on EphA2 dimerization and function.

## Acknowledgements

This work utilized the high-performance computing resources at CWRU supported by NIH grants R21AG084065, R01EY029169 and R01AG089561 to MB. We thank Dr. Paulo Cesar Telles de Souza (from University of Lyon) for communicating the new and final model of Cholesterol for Martini3 to us prior to publication. We would like to express our gratitude to Dr. Francisco Barrera for his critical reading of our manuscript.

## Author Contributions

ARS generated the protein models and performed the MD simulations and analyzed the data. MB conceived, supervised, and led the project. ARS and MB co-wrote the paper. NB provided critical advice and discussion. All authors read and approved the final version of the manuscript and supporting information.

## STAR Methods

### KEY RESOURCES TABLE

| REAGENT or RESOURCE | SOURCE | IDENTIFIER |
| --- | --- | --- |
| Deposited data |  |  |
| EphA2 TM | Bocharov et al.<sup>12</sup> | PDB ID: 2K9Y |
| Software and algorithms |  |  |
| Gromacs 2016.5/ 2021.5 | Abraham et al.<sup>46</sup> | https://www.gromacs.org/ |
| Martini 3 Force field | Souza et al.<sup>47</sup> | http://www.cgmartini.nl/ |

### RESOURCE AVAILABILITY

#### Lead contact

Further information and requests for resources should be directed to and will be fulfilled by the lead contact, Matthias Buck.

#### Materials availability

This study did not generate new unique reagents.

#### Data and code availability

This paper analyses existing, publicly available protein structures. The accession numbers for the datasets are listed in the key resources table. Molecular dynamics trajectories reported in this paper will be shared by the lead contact upon request. The paper does not report original code. Any additional information required to reanalyze the data reported in this paper is available from the lead contact upon request.

### METHOD DETAILS

#### Modeling of the TM-full JM peptides

The NMR structure of EphA2 TM dimer (PDB ID: 2K9Y)^12^ was obtained from www.rcsb.org. For modeling of the TM-JM peptide [E^530^GSGNLAV<u>IGGVAVGVVLLLVLAGVGFFI</u>HRRRKNQRARQSPEDVYFSKSEQLKPLKT YVDPHTYEDPNQAVLKFTTEIHPSC^612^], we extracted the TM region and part of the N- and C-terminal residues of EphA2 (E^530^-K^563^) from the NMR structure and then the remaining C-terminal JM residues from H^559^-C^612^ were modeled in this starting structure as an extended conformation of amino acids (ϕ, ψ= ±120°) in PyMOL (The PyMOL Molecular Graphics System, Version 2.5. Schrödinger, LLC).

#### Set up for Coarse-grain (CG) molecular dynamics simulation

To analyze the dimerization of TM-full JM peptides, the monomers were positioned 5.0 nm apart from each other. Subsequently, the atomistic (AT) models of TM-full JM peptides were transformed into a coarse-grained (CG) representation using the martinize2.py workflow module from the MARTINI 3 force field.^47^ Secondary structure assignments derived from DSSP.^48^ We employed an elastic network to enhance the stability of the helical secondary structure in the TM monomers. We used default values of the force constant of 500 kJ/mol/nm^2^ with the lower and upper elastic bond cut-off to 0.5 and 0.9 nm respectively. CG simulations were performed using GROMACS version 2016.5.64.^46^ Next, the peptides were introduced, positioned perpendicular to the membrane. We constructed two distinct systems, differentiating solely based on lipid composition: the first system comprised POPC (55%), Cholesterol (40%), and PIP2 (5%), while the second had a similar setup but without Cholesterol, consisting of POPC (95%) and PIP2 (5%). We have used the final and recently developed Martini3 CG model of cholesterol^49^ and PI(4,5)P2 model ^50^ for our systems. We utilized the insane.py^51^ script to establish the lipid bilayer around the protein. For system 1, this typically included 720 POPC, 524 CHOL, 64 PIP2 lipids, and 50181 CG water molecules. For system 2, the setup consisted of 1246 POPC, 64 PIP2, and 50110 CG water molecules. The systems were placed in a cubic box with dimensions measuring 20.0 × 20.0 × 20.0 nm^3^. The pH of the systems was considered neutral. All the simulations were run in the presence of regular MARTINI water and were neutralized to 0.15M NaCl. The systems were equilibrated for 500 ps. The long-range electrostatic interactions were used with a reaction type field having a cutoff value of 1.1 nm. We used potential-shift-verlet for the Lennard-Jones interactions with a value of 1.1 nm for the cutoff scheme and the V-rescale thermostat with a reference temperature of 320 K in combination with a Berendsen barostat with a coupling constant of 1.0 ps, compressibility of 3 × 10^-4^ bar^-1^, and a reference pressure of 1 bar was used. The integration time step was 20 fs. All the simulations were run in six replicas for 4 µs each.

#### Set up for All-Atom (AA) molecular dynamics simulation

The central conformer from each population clusters (Fig. 1C & D and Fig. S1) were selected based on the lowest RMSD value with the NMR structure of the EphA2 TM dimer (Table S1). Total of 4 dimer conformations (one from each population) were converted back to all-atom models using CG2AT2^52^ and were subjected to AA simulation with similar membrane composition as we did for the CG simulation. For system1 (+chol), both the conformers from 1^st^ and 2^nd^ population were embedded within a lipid bilayer composed of 275 POPC, 200 CHOL and 25 PIP2 molecules. For system2 (-chol), both the conformers from 1^st^ and 2^nd^ population were embedded in lipid bilayer consists of 475 POPC and 25 PIP2 molecules. All the systems were solvated using the TIP3P water model and Na/Cl ions were added to ensure charge neutrality. In addition, 150mM of NaCl was introduced using the CHARMM-GUI interface^53^ for physiological electrostatic screening. Molecular dynamics simulations were performed using GROMACS 2021.5 with the CHARMM36m force field.^54^ The long-range electrostatic interactions were managed using the particle mesh Ewald (PME) method, while hydrogen-containing covalent bonds were constrained using the SHAKE algorithm. The system temperature and pressure were controlled by Nosé-Hoover thermostat and Parrinello-Rahman type barostat. Each system underwent energy minimization through the steepest descent algorithm, consisting of 5000 steps. System equilibration followed a sequential process of six steps in CHARMM-GUI,^29^ wherein the harmonic restraints on proteins and lipids were gradually reduced. The initial two equilibration steps were conducted within the NVT ensemble, while the subsequent four steps occurred under the NPT ensemble to maintain an isobaric-isothermal condition. A total of four different simulations were conducted (two for each system) for 200ns each and all the simulations were run in quadruplicates.

#### Quantification and statistical Analysis

Trajectory analysis was conducted using the integrated modules within GROMACS. Contact maps depicting the TM regions and with the lipids were generated, employing a cutoff of 6 Å for both backbone and side-chain non-hydrogen atoms. Subsequently, the data was visualized and plotted using Origin.^55^ Protein-lipid interactions were analyzed using vmd^56^ and PyLipID.^57^ Local membrane properties are analyzed using the *g_lomepro* program.^58^ The inter-helical angle and distance were calculated as described elsewhere.^37^ We have used PyMOL^59^ and ChimeraX ^60^ for preparing the figures.

#### Supporting Information Available

Addition figures are included in the supporting information.

