## supplementary file for "Cholesterol-Dependent Dimerization and Conformational Dynamics of EphA2 Receptors: Insights from Coarse-Grained and All-Atom Simulations"

Supplemental Table S1 and Figures S1 to S4 are provided.

**Table S1:** Comparison of CG simulated EphA2 TM dimers with the NMR structures. Central conformers of the populated clusters from the CG simulation are considered for calculating the mean and SD of the crossing angle values.

| Membrane | CG simulation | | | | NMR |
|---|---|---|---|---|---|
| | X (deg) | | RMSD (Å) from NMR | | X (deg) |
| | 1st Cluster | 2nd Cluster | 1st Cluster | 2nd Cluster | |
| +CHOL | 35.6 ± 6.4 | -29.1 ± 6.2 | 4.2 ± 0.4 | 4.5 ± 0.1 | 17 |
| -CHOL | 25.0 ± 6.1 | 7.0 ± 3.8 | 5.1 ± 0.2 | 4.4 ± 0.4 | |

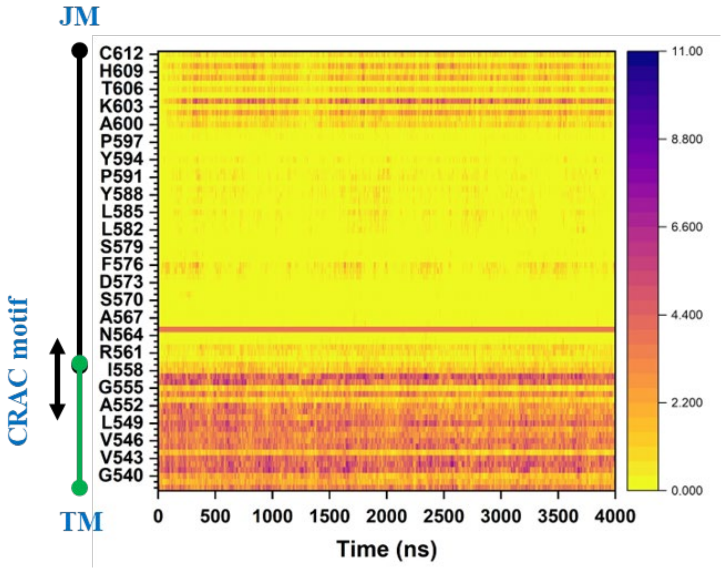


**Fig. S1** Comparison of the average number of interactions between the TM-full JM peptides with Cholesterol molecules over the course of CG simulation. The TM and the JM region are marked on the Y- axis scale. The region of the CRAC motif (V-X1-5-F-X1-5-R) on the TM domain are also marked. We used 0.7 nm cutoff for this calculation.

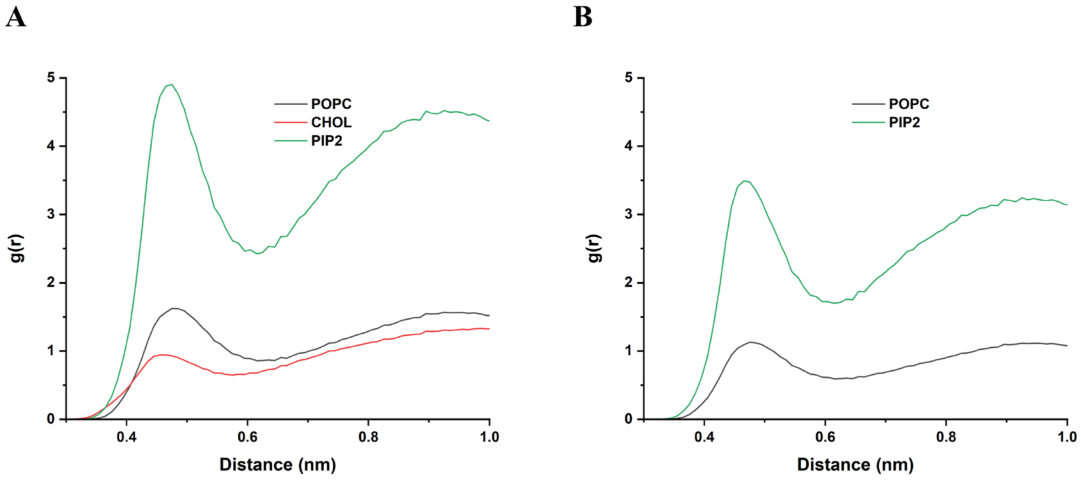


**Fig. S2** Comparison of the radial distribution functions of lipids around the protein for both systems: (A) with Cholesterol and (B) without Cholesterol from the CG simulations. All the values converge at ~1 at longer distances (not shown here).

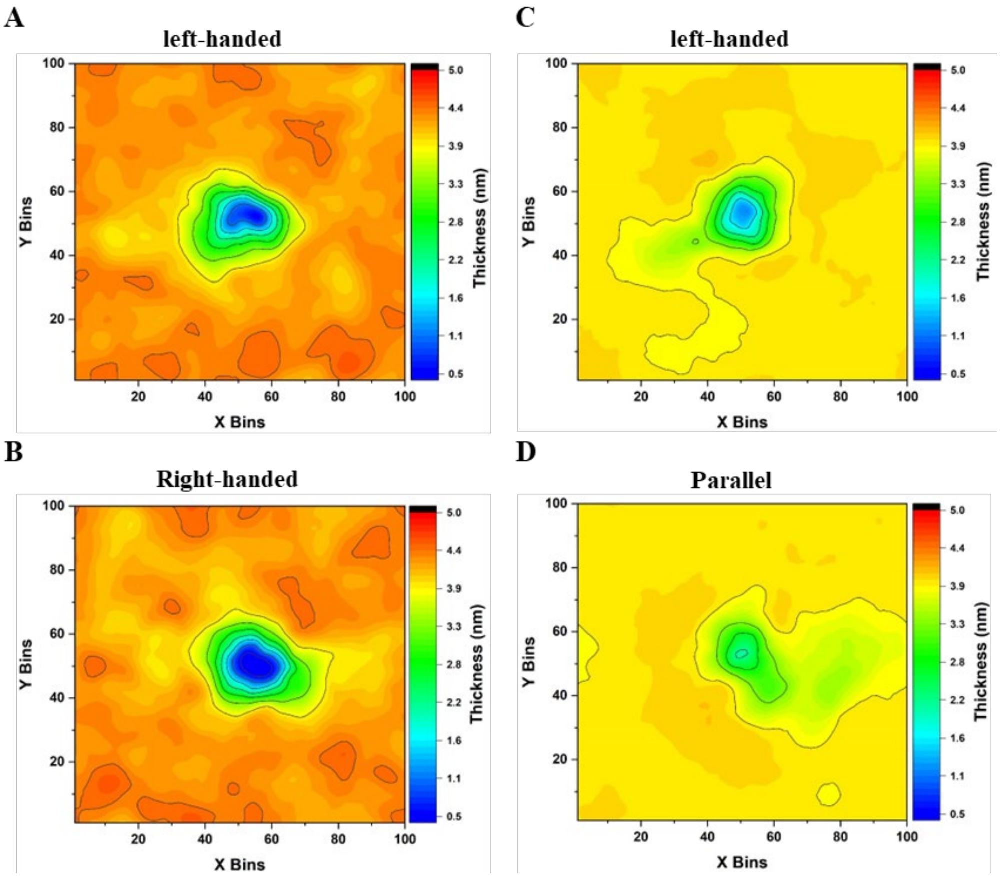


**Fig. S3** Comparison of local membrane thickness calculated over the entire membrane for the Cholesterol rich (A & B) and cholesterol deficient (C & D) systems from the AA simulations. Replica trajectories are combined for this analysis. X and Y bins refer to grid dimensions (0.1 by 0.1 nm) across the membrane plane (xy plane).

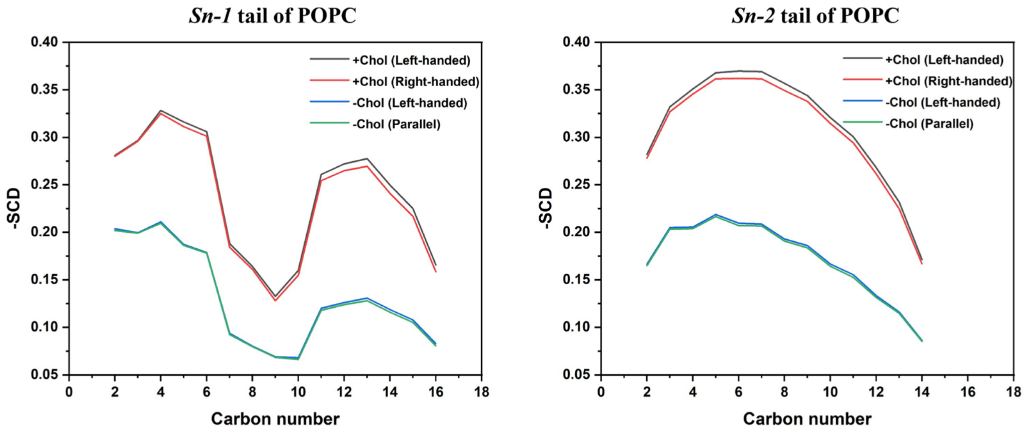


**Fig. S4** SCD comparison of both sn-1 and sn-2 tail for POPC lipids in cholesterol-rich and cholesterol-deficient systems from the AA simulations. Replica trajectories are combined for this analysis.
